# Geometric, cell cycle and maternal-to-zygotic transition-associated YAP dynamics during preimplantation embryo development

**DOI:** 10.1101/2025.02.27.640568

**Authors:** Madeleine Chalifoux, Maria Avdeeva, Eszter Posfai

## Abstract

During the first cell fate decision in mammalian embryos the inner cell mass cells, which will give rise to the embryo proper and other extraembryonic tissues, segregate from the trophectoderm cells, the precursors of the placenta. Cell fate segregation proceeds in a gradual manner encompassing two rounds of cell division, as well as cell positional and morphological changes. While it is known that the activity of the Hippo signaling pathway and the subcellular localization of its downstream effector YAP dictate lineage specific gene expression, the response of YAP to these dynamic cellular changes remains incompletely understood. Here we address these questions by quantitative live imaging of endogenously tagged YAP while simultaneously monitoring geometric cellular features and cell cycle progression throughout cell fate segregation. We apply a probabilistic model to our dynamic data, providing a quantitative characterization of the mutual effects of YAP and cellular relative exposed area, which has previously been shown to correlate with subcellular YAP localization in fixed samples. Additionally, we study how nuclear YAP levels are influenced by other factors, such as the decreasing pool of maternally provided YAP that is partitioned to daughter cells through cleavage divisions, cell cycle-associated nuclear volume changes, and a delay after divisions in adjusting YAP levels to new cell positions. Interestingly, we find that establishing low nuclear YAP levels required for the inner cell mass fate is largely achieved by passive cell cycle-associated mechanisms. Moreover, contrary to expectations, we find that mechanical perturbations that result in cell shape changes do not influence YAP localization in the embryo. Together our work identifies how various inputs are integrated over a dynamic developmental time course to shape the levels of a key molecular determinant of the first cell fate choice.

## Introduction

During embryonic development cells use cues from their environment and/or asymmetrically inherited factors to instruct fate specific gene expression. Cell fate acquisition is typically not instantaneous, but rather a gradual process that takes place while cells progress through cell cycles, change geometric properties such as shape and position, and transit from maternal to zygotic resources - all of which can impinge on cell fate establishment. To address how these dynamic inputs are integrated and how they shape the levels of key cell fate effectors, quantitative live imaging approaches are needed to simultaneously measure relevant cellular features along with fate determinants. Here, using the first cell fate decision in the mouse embryo as an example, we investigate how dynamic inputs collectively affect the Hippo signaling effector Yes-associated protein (YAP), a key determinant of this fate choice.

In mammals, the first two cell types to differentiate are the trophectoderm (TE) and the inner cell mass (ICM), setting aside the precursors of the placenta from the cells which will go on to give rise to the future fetus, respectively. By the blastocyst stage both cell types are well defined, with the TE forming a single epithelial layer on the surface, and the ICM cells allocated to the inside of the embryo. A key factor regulating this early fate decision is the transcriptional co-factor YAP, which when localized to the nucleus interacts with the transcription factor TEAD4 to activate *Cdx2* and repress *Sox2* expression, regulators of the TE and ICM, respectively [1], [2], [3], [4], [5]. Subcellular YAP localization in turn is dictated by the Hippo signaling pathway [3]. In ICM cells, Hippo signaling is active and YAP is phosphorylated by the LATS 1/2 kinases resulting in its cytoplasmic localization [3], [6]. In TE cells, on the other hand, Hippo signaling is inactive, and unphosphorylated YAP localizes to the nucleus.

A well-established regulator of the Hippo pathway in preimplantation embryos is apico-basal polarity. In apolar cells, destined to become ICM, the Hippo pathway member angiomotin (AMOT) is localized to adherens junctions, where it activates the LATS1/2 kinases [7], [8]. In contrast, TE cells positioned to the surface of the embryo establish distinct membrane domains: basolateral polarity proteins segregate to cell-cell contact surfaces, while exposed surfaces accumulate apical polarity proteins and establish an apical domain (AD) [9], [10], [11], [12]. The AD sequesters AMOT, preventing it from engaging in Hippo signaling [7], [8]. Interestingly, the proportion of exposed cell surface area has been shown to correlate with subcellular YAP localization, suggesting its potential role as a continuous readout of a cell’s polarity status [13].

While the role of polarity/REA in regulating YAP localization in the early embryo is well-appreciated, YAP has been shown to respond to mechanical forces in a variety of other systems [14], [15]. Mechanosensitivity of YAP in the embryo has also been put forward, specifically, response of YAP to cortical contractility (or cortical tension) has been proposed [16], [17]. However, the relationship between YAP localization and the degree of contractility is not linear, e.g. cells with the lowest and highest amount of contractility display cytoplasmic YAP localization, while cells with intermediate contractility localize YAP to the nucleus. One possibility is that YAP localization is rather sensitive to cell shape deformations [18]. Indeed, in cells that are more round (low aspect ratio) YAP is cytoplasmic despite variable contractility states, while outer flattened cells (high aspect ratio) localize YAP to the nucleus [3], [13], [19]. Cell shape deformations can arise due to asymmetries in the contractility states of neighboring cells or from the forces stemming from the formation of the blastocoel cavity [16], [20], [21]. While long-term shape deformations have been applied to developing embryos, these perturbations also altered REA [13] and thus likely polarity, therefore a direct role of cell shape on YAP localization remains unknown.

Here, we use a live reporter for endogenous YAP (YAP-miRFP670) [22] to directly visualize and track its dynamics in individual cell lineages from the 8- to the 32-cell stages of development, during the segregation of ICM and TE. We reveal how the concerted effects of transitioning from maternal to zygotic YAP, cell cycle dynamics and cell positional but not cell shape changes influence YAP dynamics. Together, these inputs set the diverging YAP dynamics in cells that will form ICM or TE.

## Results

### Live imaging nuclear YAP dynamics during inner cell mass and trophectoderm segregation

To visualize nuclear YAP dynamics throughout the first cell fate decision, we used the *Yap-miRFP670* reporter mouse line, in which endogenous *Yap* is fluorescently tagged [22]. Homozygous *Yap-miRFP670* embryos were derived by crossing homozygous female and male animals (Fig. 1A). To simultaneously visualize nuclei, embryos were isolated at the 2-cell stage and injected with *H2B-miRFP720* mRNA. Embryos were then live imaged from the 8- (embryonic day (E)2.5) through 32-cell stage (∼E3.5) using an Inverted Lightsheet Microscope (InVi) [23] acquiring a z-stack every 15 minutes for ∼30 hours (Fig. 1B). Using our previously established semi-automatic image analysis pipeline, nuclei were segmented, 3D nuclear masks were registered between time frames and tracked through time to construct cell lineages (Fig. 1C, Fig. S1A, B, and C) [24]. Nuclear YAP fluorescent signal intensity was extracted from each nucleus at each point in time, allowing us to analyze nuclear YAP concentration (total YAP fluorescent intensity normalized by nuclear volume) dynamics in individual cell lineages, starting from a single 8-cell stage blastomere to its four progeny at the 32-cell stage (Methods, Fig. 1D). These data allow us to assess the dynamics of YAP behavior throughout the segregation of the ICM and TE lineages as cells progress through cell cycles, and positional changes and shape deformations.

**Figure 1.**
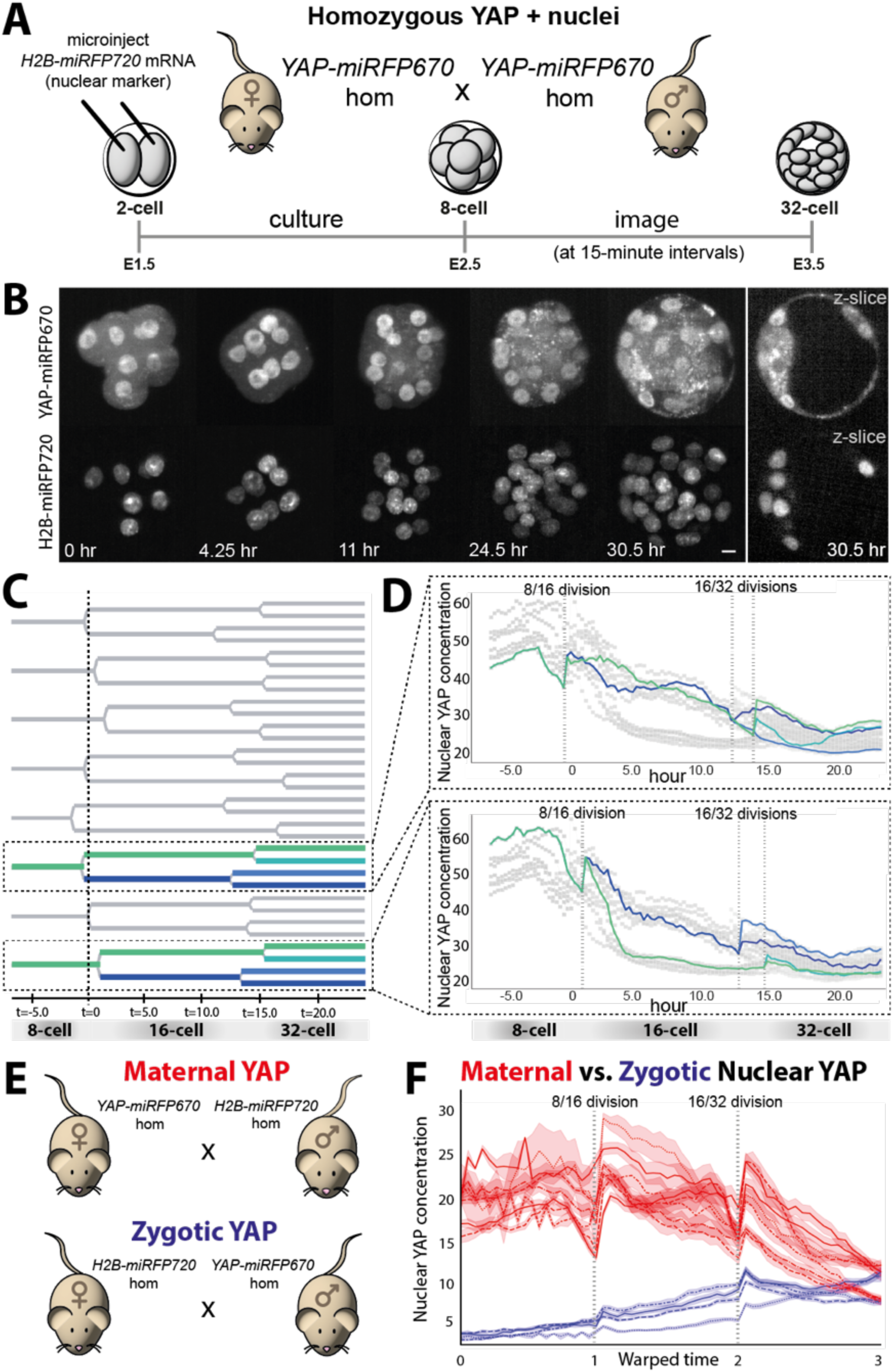
Long-term live imaging of preimplantation embryos expressing endogenously-tagged YAP-miRFP670. A. Schematic illustrating experimental genetics and setup: Female and male mice homozygous (hom) for *YAP-miRFP670* are crossed. E1.5 (2-cell) embryos derived from this cross are microinjected with *H2B-miRFP720* mRNA as a nuclear marker and cultured until E2.5 (8-cell). Embryos are imaged on a light sheet microscope at 15-minute intervals from the 8- through 32-cell stage. B. Time lapse images of a YAP-miRFP670-expressing embryo imaged alongside labeled nuclei (H2B-miRFP670) from the 8-cell through early blastocyst stage. A z-stack was acquired for both channels every 15 minutes. Maximum intensity projections (MIPs) are shown, unless otherwise specified as a single plane z-slice. Unit of time is hours. Scale bar is 10um. C. Example lineage tree for a YAP-miRFP670-expressing embryo from the 8- through the 32-cell stage, as marked by the time point preceding the first division into 64- cell stage. Highlighted branches correspond to the traces displayed in panel D. t=0 represents the average time of division from the 8-16 cell stage within the whole embryo. Unit of time is hours. D. Example traces of lineages (extracted from two branches of the lineage tree in C) showing the dynamics of nuclear YAP concentration (average reporter intensity in the nucleus). Each trace follows an 8-cell stage blastomere that divides to two daughters at the 16-cell stage, followed by division to four granddaughters at the 32-cell stage. t=0 represents the average time of division from the 8-16 cell stage within the whole embryo. Unit of time is hours. E. Schematic illustrating genetics of crosses to obtain embryos with maternal (reports on maternal contribution and one zygotic allele) or zygotic (reports on one zygotic allele) YAP-miRFP670. Maternal YAP crosses are obtained by mating a homozygous *YAP-miRFP670* female to a male homozygous for *H2B-miRFP720* (nuclear marker). Zygotic YAP crosses are obtained by mating a homozygous *H2B- miRFP720* female to a male homozygous for *YAP-miRFP670*. F. Quantification of nuclear YAP concentration over time in embryos with maternal or zygotic YAP-miRFP670, imaged on a light sheet microscope. Each line is the average of all traces in an embryo, shaded area represents the 95% confidence interval. Warped time (see Methods) is aligned by 8/16 and 16/32 divisions. N=7 maternal embryos, N=4 zygotic embryos.

### Distinct maternal and zygotic YAP contributions

A striking feature of YAP dynamics we observed was the gradual decline in YAP concentration in all cells during development. Both *Yap* mRNA and YAP protein are known to be present in the oocyte [25], therefore we hypothesized that the gradual decrease reflected depletion of the maternal pool of YAP. To test this, we generated embryos in which *Yap-miRFP670* was either inherited maternally (reports on maternal pool of YAP and the maternally inherited zygotic *Yap* allele) or paternally (reports on the paternally inherited zygotic *Yap* allele) (Fig.1E). Live imaging nuclear YAP in these groups of embryos showed that indeed, a larger and relatively stable pool of maternal YAP is present up to the 32-cell stage, while lower levels of YAP originating from the paternally inherited zygotic allele are first evident only at the 16-cell stage (Fig. 1F and Fig. S1D). Only by the mid-32 cell stage do we find similar nuclear YAP levels from maternal and paternal sources, indicating that early fate patterning events in the embryo largely use YAP of maternal origin.

### YAP dynamics through cell cycles

In agreement with previous work [22], we also observed “spikes” of nuclear YAP concentration increase following cell divisions (both 8/16 and 16/32 divisions) which were evident in nearly all lineages (Fig. 1D and 1F). To understand the origins of these spikes, we investigated how YAP dynamics are affected by cell division. Specifically, we analyzed how the total amount of nuclear YAP is inherited through divisions, using the 8/16-cell division as an example. Averaging total nuclear YAP in the hour before nuclear envelope breakdown and in the hour after nuclear envelope re-establishment (Fig. S2A), respectively, we found that the pool of un-phosphorylated YAP from the mother cell is losslessly inherited by the two daughter cells (Fig. 2A) and is initially partitioned into the nuclei of the two daughters at similar levels (Fig. 2B).

**Figure 2.**
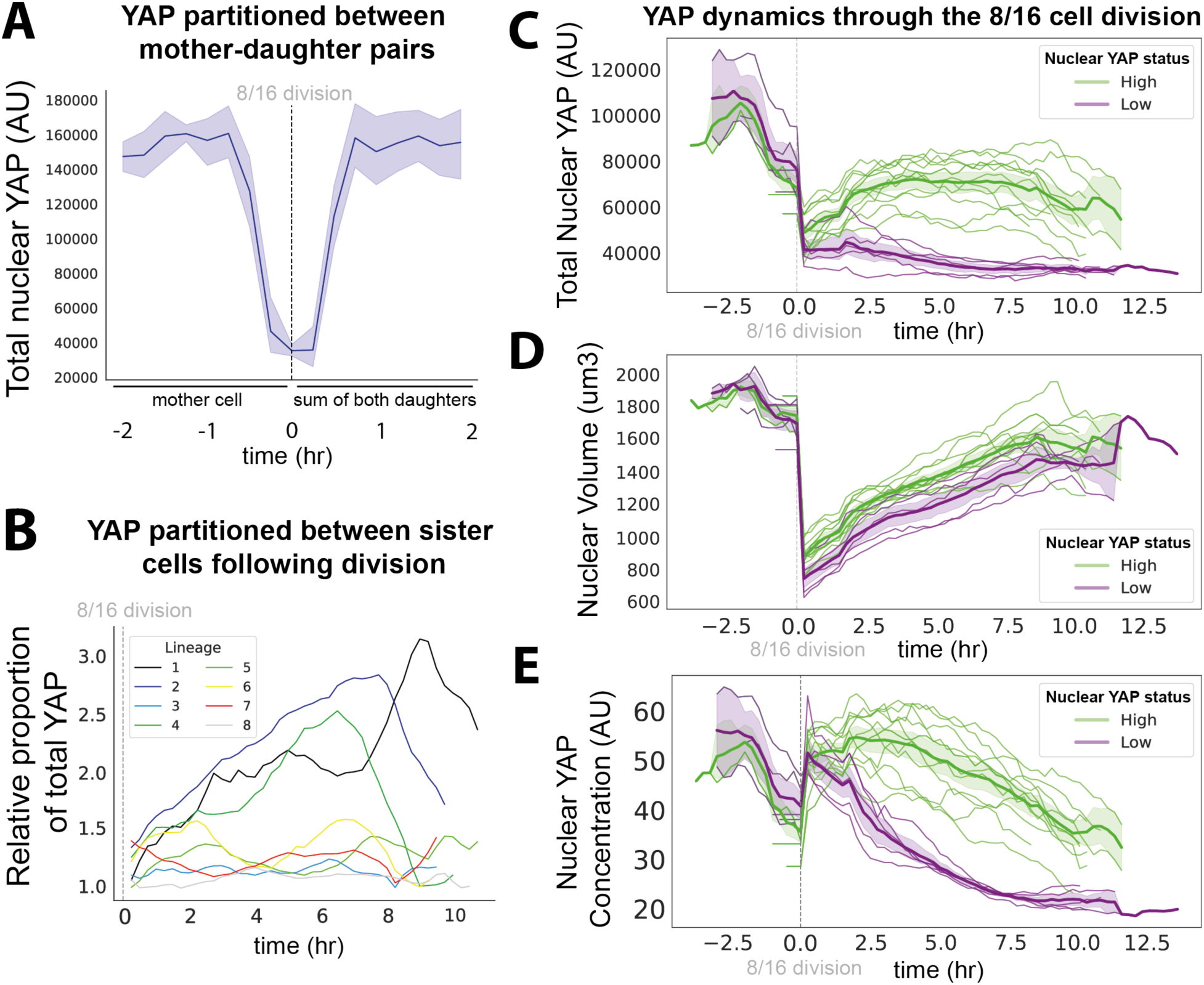
YAP dynamics during cell cycles. A. Total nuclear YAP (sum of reporter intensity in the nucleus) measured in the 8-cell stage mother cell and the total amount of nuclear YAP contained within its two daughter cells at the 16-cell stage. n=8 mother cells and 16 daughters within one embryo. Solid blue line shows average, shaded blue represents the 95% confidence interval. Quantification is representative of all embryos (N=12) analyzed. Traces are aligned to the 8/16 cell division, shown as t=0. Unit of time is hours. AU - arbitrary units. B. Traces showing the relative proportion (larger over smaller total nuclear YAP values) of total nuclear YAP inherited by sister cell pairs starting at the 8/16 division (t=0) from one embryo. Unit of time is hours. Quantification is representative of all embryos (N=12) analyzed. C. Traces showing total nuclear YAP (AU) in individual lineages from an embryo (data representative of N=13 embryos). Traces are aligned to 8/16 cell division (dashed vertical grey line). Unit of time is hours. Cell lineages are classified into high nuclear YAP (green) and low nuclear YAP (purple) populations based on their late 16-cell stage YAP status (see Methods). Each fine green and purple line corresponds to a cell lineage, solid green and purple lines show averages over the corresponding subpopulations, shaded green and purple regions show the 95% confidence interval. D. Same as (C) showing nuclear volumes (um^3^). E. Same as (C) showing nuclear YAP concentration (AU).

To assay how YAP levels diverge among daughters following division, we first annotated lineages at the 16-cell stage by their YAP status. We found that the distribution of average nuclear YAP expression at the late 16-cell stage had two major modes (Fig. S3B, Methods), which allowed us to classify cells as low or high YAP (Fig. 2C, D and E). Using this annotation, we aligned lineages at the 8/16-cell division and observed a gradual divergence of total YAP levels between the low and high YAP branches, reaching a steady state approximately 2-3 hours after division (Fig. 2C). This indicates that several hours are likely necessary for Hippo signaling differences to adjust YAP levels according to the newly acquired position and polarity status of the cell. Notably, we found that most change in YAP levels occurred in the high YAP branches, indicating that additional YAP enters the nucleus, presumably due to de-phosphorylation of the cytoplasmic YAP pool. In contrast, in YAP low branches the decrease of total nuclear YAP after division was minimal, suggesting a minor contribution of nuclear YAP export in setting the low YAP subpopulation.

After determining the behavior of total YAP levels, we turned to analyzing how nuclear volume dynamics affect the nuclear concentration of YAP. We found a drastic reduction of nuclear volume immediately after division, followed by a decelerating expansion of the volume (Fig. 2D). Taking these dynamics into account, we find they result in prominent spikes in nuclear YAP concentration after divisions in both YAP high and low lineages (Fig. 2E).

In summary, we find that at divisions the total amount of nuclear YAP is losslessly partitioned into daughter cells at similar levels. Within 2-3 hours after division, low and high YAP subpopulations diverge to adjust to new position and polarity states, which mainly manifests by outer cells importing additional YAP into the nucleus. In addition, changes in nuclear volumes also significantly affect nuclear YAP concentration. Specifically, initial nuclear volume reduction causes YAP concentration spikes, followed by nuclear volume expansion lowering YAP concentration. These measurements provide novel insight into the effects cell cycle dynamics have on nuclear YAP levels during the first cell fate decision and in particular, reveal that emergence of the low YAP state is largely due to dilution of nuclear YAP through cleavage division and nuclear volume expansion.

### Nuclear YAP levels dynamically establish a strong correlation with Relative Exposed Area

Previous work in fixed embryos identified a strong correlation between a cell’s REA and nuclear/cytoplasmic YAP ratio across the late 8- to 64-cell stages [13]. To assay the dynamics of this correlation during developmental cell cycles, we live imaged embryos expressing H2B-miRFP720, YAP-miRFP670, as well as membraneTomato (using the *mTmG* mouse line [26] (Fig. 3A). To measure the relative exposed area (REA) of each cell at each time point, we used Cellpose to segment cell membranes and calculated the membrane area that was or was not in contact with any other cell membrane [27] (Fig. 3B, Methods). We define REA as the area of the contact-free surface divided by the entire surface area of the cell (Fig. 3C). We then correlated REA with nuclear YAP levels separately throughout the 16- and 32-cell stages, by aligning each cell lineage to the time of the 8/16 or 16/32 cell division, respectively (Fig. 3D). Overall, for both the 16- and 32- cell stages we observed a weaker Pearson correlation at the beginning of each cell cycle, that gradually increased within the first 2-3 hours reaching an overall strong correlation for the remainder of the cycle. This is in line with our previous observation that several hours are needed to adjust YAP levels to new cell positions and polarity states acquired after cell division and shows that following this adjustment YAP better reflects Hippo activity determined by the relative size of these different membrane domains. Overall, we noted that the REA:YAP correlation is time-dependent within and between cell cycles which prompted us to further model their relationship with dynamic Bayesian networks (DBNs).

**Figure 3.**
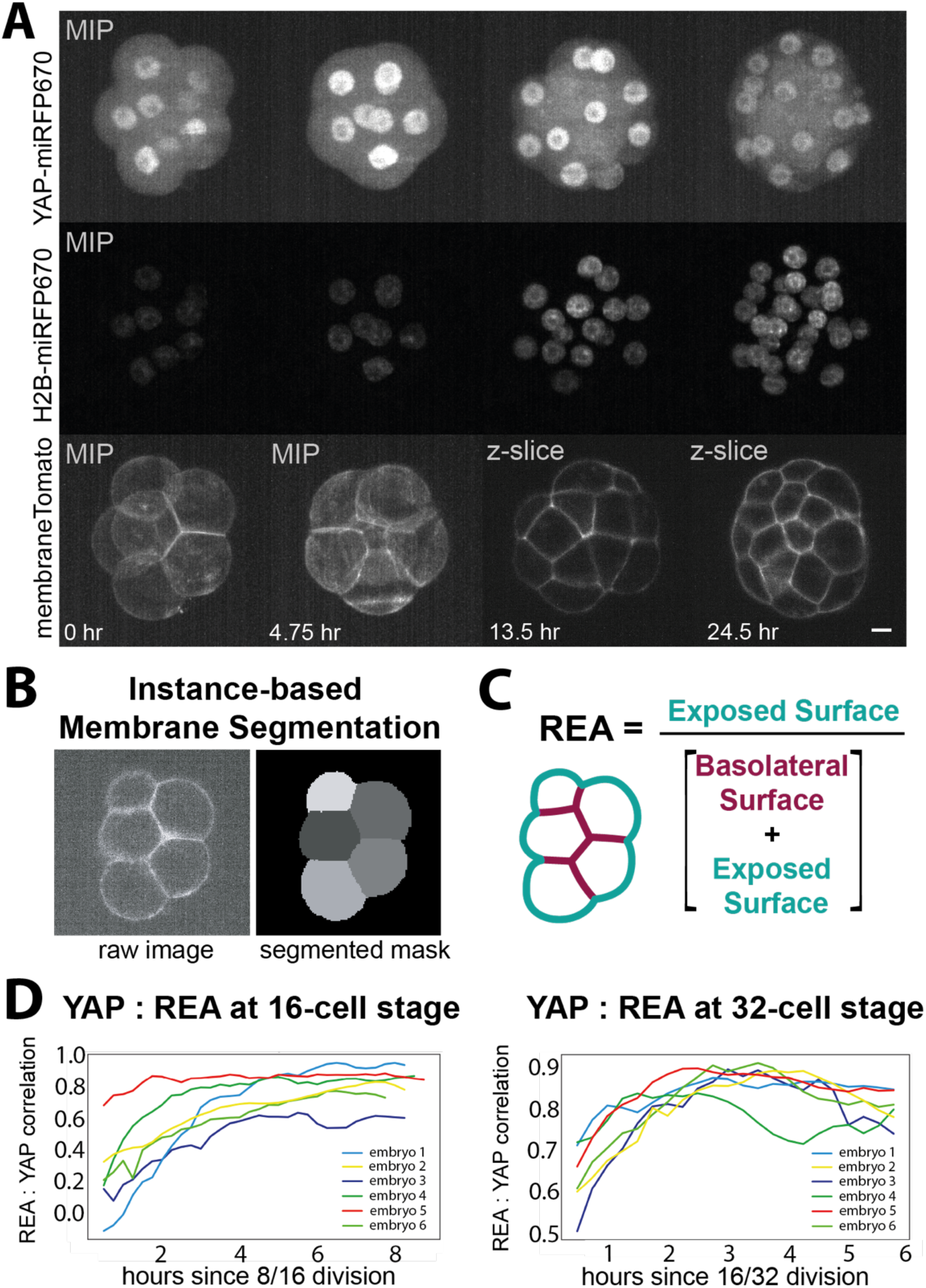
Establishing correlation between YAP and REA during development. A. Time lapse imaging of an embryo expressing YAP-miRFP670 alongside nuclear (H2B-miRFP720) and membrane (membraneTomato) markers from the 8- to 32- cell stages. A z-stack was acquired in all three channels every 15 minutes. Images shown are maximum intensity projections (MIPs), except for the membrane images at t=13.5 and t=24.5 hours, which are of a single z-plane (z-slices). Unit of time is hours. Scale bar is 10um. B. An example membrane image (z-slice) and its associated membrane mask automatically generated using an instance-based membrane segmentation algorithm (Cellpose). C. Relative exposed area (REA) is defined by a cell’s exposed surface area normalized by the total cell surface area. Pearson correlation coefficient between nuclear YAP concentration and REA observations across all cells, calculated at each time point during the 16- (left) and 32-cell (right) stages. Color: embryo. N=6 embryos. Unit of time: hours.

### Modeling YAP and Relative Exposed Area dynamics

To extract the salient patterns in the relationship between REA and YAP localization dynamics and interpret the corresponding parameters, we turned to Bayesian techniques. To this end, we assumed a DBN approach which we have previously successfully applied to model the dynamics of the first cell fate specification in the early embryo [28].Briefly, we model the data using a graph with categorical random variables in the nodes, corresponding to discretized levels of REA or nuclear YAP over 5 coarse-grained timepoints, 8-cell (0), early (1) and late (2) 16-cell and early (3) and late (4) 32- cell stages (Methods). While the levels of YAP were discretized as high (1) and low (0), we classified cells as inner (REA=0), intermediate (REA=1), and outer (REA=2) (Methods, Fig. S3A and B). In DBNs, the network is endowed with additional structure where the nodes are indexed by time, describing the state of the system at a particular timepoint. Edges in a Bayesian network describe dependencies in the data (Methods), and in DBNs, only edges between the nodes in the same or adjacent time slices are considered, with the parents belonging to the same or the previous time slice. In this work, we considered 10 possible networks describing the potential YAP-REA relationship (Fig. S3C). The networks were assessed via their structure score calculated using Bayesian Information Criterion (Methods, Fig. S3D), and the network with the highest score was selected to further model the data. The selected network incorporates fast effects of REA on YAP localization (R_t → Y_t edges), with upstream dynamics of REA governing dynamics of YAP (Fig. 4A). Interestingly, the structure also includes an additional edge with the slower feedback of YAP into REA at the 16-cell stage (Y_1 → R_2 edge).

**Figure 4.**
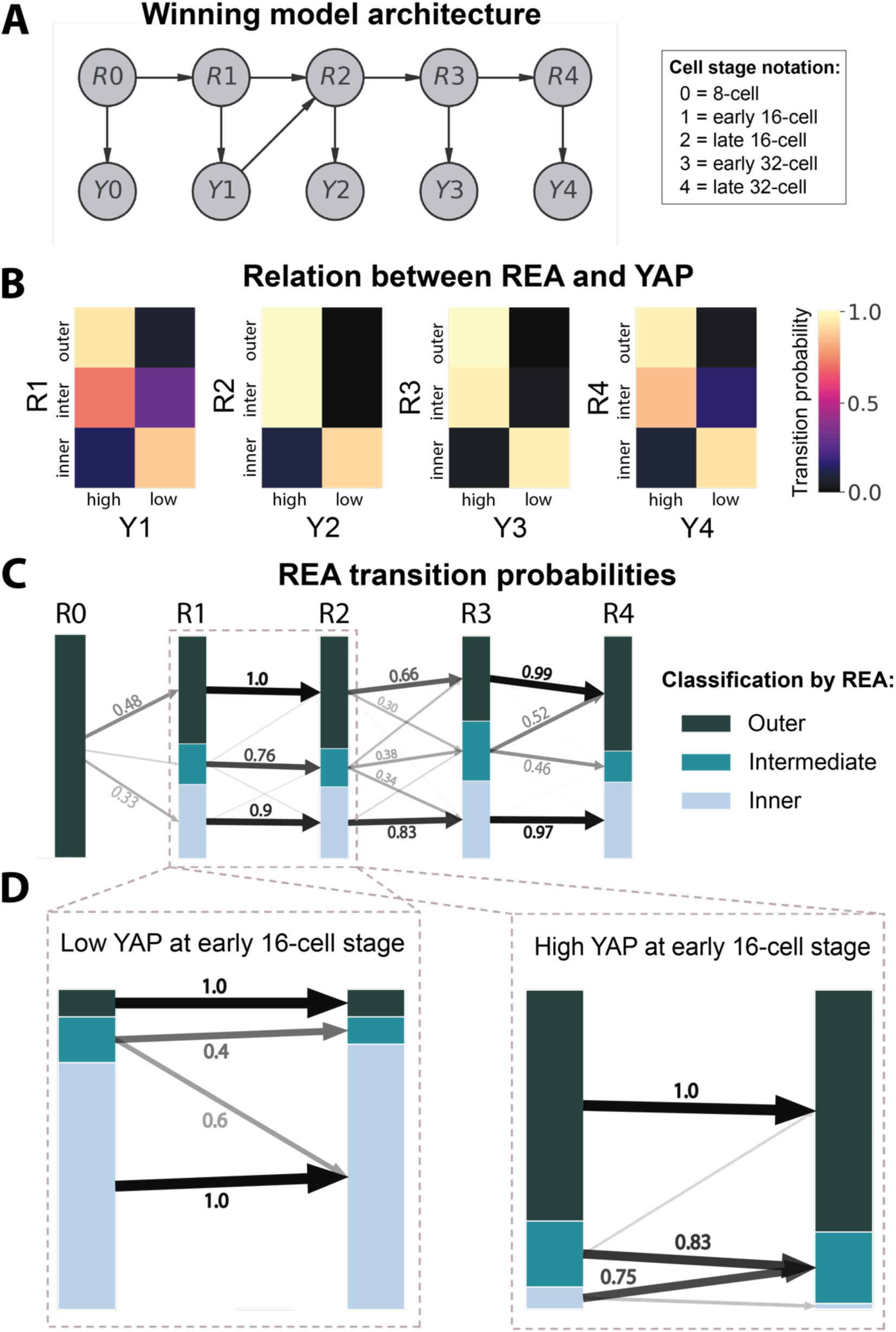
Modeling the relationship between YAP and REA. A. A dynamic Bayesian network applied to model REA-YAP dynamics. R_t, Y_t, t=0…4: discretized REA and YAP levels summarized over 5 stages, modeled via categorical distributions. Edges encode dependencies in the data (See Methods). B. Conditional probability distributions for R_t -> Y_t edges, shown for early and late 16-cell stages, as well as for early and late 32-cell stages. For every stage, REA can assume three states while YAP is binary. Probability is shown in color. C. Conditional probability distributions for R_t -> R_{t+1} edges. Stacked barplots demonstrate distribution of REA at every stage, bar height is proportional to state probability. Arrows: transition probabilities. For every two states at adjacent stages, the arrows connect the corresponding bars; thickness and transparency are proportional to the probability. All arrows with probability >0.3 are labeled. For the CPD between stages 1 and 2, Y_1 was marginalized out. D. R_1 -> R_2 CPDs conditioned on Y_1=0 (left) and Y_1=1 (right). Barplots and arrows: same as (C).

For every edge of the network, a conditional probability distribution (CPD) describes the corresponding dependencies. In particular, we learned the structure of R → Y transition probability matrices for every timepoint (Fig. 4B). While at the early 16- cell stage, the matrix contains some non-deterministic elements, with intermediate cells demonstrating variable YAP behavior, at later stages, inner cells display low nuclear YAP while intermediate and outer cells have high nuclear YAP with overwhelming probability. This is consistent with previous observations that a cell’s position (e.g. REA) at the early 16-cell stage is not fully representative of its polarity state, and that YAP rather responds to the polarity state of cells than REA [20], [29]. As from the late 16-cell stage onwards REA reflects a cell’s polarity state [30], [31], REA is a strong predictor of YAP levels at these stages.

With a simple structure of the network and low stochasticity observed in R → Y CPDs, the dynamics of YAP can be easily described in terms of REA dynamics shown in Fig. 4C. Outer cells consistently form the largest subpopulation; intermediate cells, on the other hand, are the smallest subpopulation which appears at the early 16-cell stage and persists through late 32-cell stage. REA is mostly conserved during interphases (R_1 → R_2 and R_3 → R_4) and is most likely to be perturbed at divisions (R_0 → R_1 and R_2 → R_3).

In particular, while during the 16-cell interphase REA is indeed most likely to be conserved, its dynamics is dependent on the YAP status of the cell, with transition probabilities significantly different between low and high YAP cells (Fig. 4D). The differences between these transition probabilities are most pronounced in the intermediate REA subpopulation. Low YAP intermediate REA cells at the early 16-cell stage are likely to be internalized, which is consistent with previous observations of internalization of apolar cells with an exposed surface area [16], [32]. At the same time high YAP intermediate cells are most likely to stay intermediate or externalize even further to become outer cells. In summary, we find that YAP status at the early 16-cell stage is predictive of a cells position dynamics until the next division.

Our modeling approach provides a quantitative overview of the dynamic relationship between positional information and YAP, highlighting the extent this geometric feature can explain nuclear YAP levels.

### YAP localization does not respond to cell shape deformations

While it is evident that a cell’s polarity state (approximated by REA) regulates YAP localization, we asked whether additional inputs, such as mechanical forces that result in cell shape deformations could also contribute to YAP regulation.

We performed mechanical perturbations on time scales (up to 90 minutes) that are not expected to affect polarization of most cells but have been shown to alter YAP localization in different cellular systems [18], [33], [34], [35], [36], [37]. First, we compressed 16-cell stage embryos expressing YAP-miRFP670 to a height of 30 μm, which deformed the shape of blastomeres (Fig. 5A and B). We confirmed that even after 90 minutes this perturbation either did not result in the exposure of inner cells or, if a small exposed surface area did appear, it did not polarize Ezrin, a component of the AD (Fig. S4A). By measuring the aspect ratio of cells before and after compression, we found that the aspect ratio increased, resembling aspect ratios of E3.5 TE cells (Fig. S4B and C). If YAP is sensitive to such mechanical perturbation, we would expect an increase in nuclear YAP levels. We acquired an image of each embryo prior to compression, and for a duration of 90 minutes post-compression, imaging every 10 minutes (Fig. 5B). Nuclei were segmented and YAP-miRFP670 fluorescent intensity was extracted and normalized by histone intensity to account for changes in distance to the objective pre- and post-compression. We found that nuclear YAP-miRFP670 levels did not increase during compression, but instead showed a slight decrease over time, consistent with degradation of maternal protein (N=9 embryos, n=150 cells) (Fig. 5C and Fig. S4D). We also tracked individual nuclei through 10-90 minutes of manipulation and observed no increase in nuclear YAP levels (Fig. S4E).

**Figure 5.**
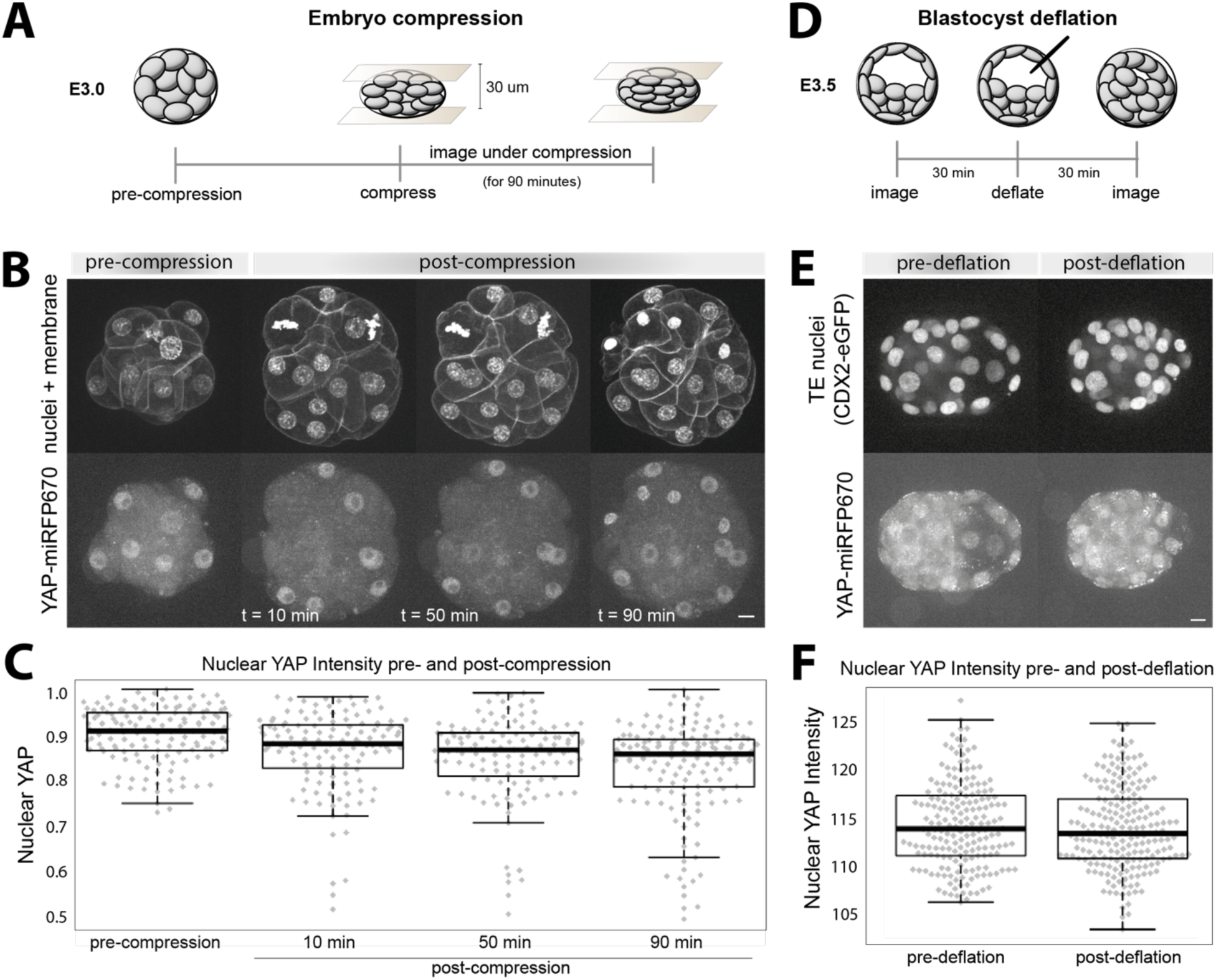
YAP localization does not respond to two types of mechanical perturbations. A. Schematic of embryo compression. Late 16- or early 32-cell stage embryos were used (E3.0). An initial pre-compression z-stack is acquired for each embryo, followed by compressing the embryo to a height of 30um. Embryos were imaged for 90 minutes under compression, acquiring a z-stack every 10 minutes. B. Example images of YAP-miRFP670, H2B-miRFP720 and membraneTomato-expressing embryos subjected to compression. Time indicates minutes after compression. Scale bar 10um. C. Nuclear YAP Intensity (AU) normalized by histone intensity (AU) in individual cells, quantified pre-and post compression for all embryos subjected to compression (N=9 embryos, n=164 cells). Cells actively undergoing mitosis were excluded from the analysis. D. Schematic of blastocyst deflation. E3.5 blastocysts just beginning cavitation were imaged before and ∼30 minutes after blastocoel deflation. Deflation was performed using a fine glass capillary. E. Example images of YAP-miRFP670, CDX2-eGFP-expressing blastocysts subjected to deflation. An initial pre-deflation z-stack was acquired for each embryo, followed by a post-deflation z-stack 30 minutes after blastocoel deflation. CDX2-eGFP selectively labels TE cells. Scale bar 10um. Quantification of nuclear YAP intensity (AU) in individual TE cells of all pre- and post-deflation blastocysts analyzed (N=6 embryos, n=202 cells).

Embryo compression demonstrated that cells with direct force application and increased cell aspect ratio do not result in higher levels of nuclear YAP within the time period imaged. Conversely, to assess how decreased aspect ratio affects YAP localization, we deflated blastocoel cavities of E3.5 blastocysts (Fig. 5D and E). During normal development, inflation of the blastocoel cavity causes TE cells to stretch and thus increase in aspect ratio [21]. Deflation relieves pressure in the blastocoel, causing TE cells to round up and their aspect ratio to decrease (Fig. S4F). We acquired images of YAP-miRFP670 embryos also expressing a TE marker (CDX2-eGFP) before and 30 minutes after blastocoel deflation (Fig. 5E). Quantification of nuclear YAP in TE cells of all embryos (N=6 embryos, n=202 cells) demonstrated that nuclear YAP levels did not significantly change before and after deflation (Fig. 5F). Therefore, in contrast to expectations, we find that these different cell shape deformations do not affect YAP localization in the preimplantation embryo.

## Discussion

Cell fates during early embryonic development are acquired concomitant with transitions from maternal to zygotic gene products, dynamic morphological changes to cell position and shape and cell cycles. Here by using live imaging we investigate how blastomeres of the preimplantation embryo integrate such cues to regulate the levels and subcellular localization of YAP, a key determinant for the ICM/TE fate choice.

Until now, studies have focused on analyzing YAP nuclear:cytoplasmic localization as the primary readout of YAP behavior [3], [13], [38]. However, with the use of quantitative live imaging, we are able to investigate the amount of YAP during preimplantation development. We show that a large pool of maternally supplied YAP is present in the early embryo that gradually decreases until the 32-cell stage, while zygotically expressed YAP is first weakly detected by the 16-cell stage. Thus, early cell fate dynamics largely utilize YAP of maternal origin, highlighting the importance of maternally deposited YAP in the oocyte. These observations are underscored by previous work demonstrating that loss of maternal YAP led to precautious activation if the ICM marker SOX2 [39].

ICM fate requires low nuclear YAP concentration. While it is well-appreciated that lack of cell polarization and therefore Hippo signaling activation is needed for low nuclear YAP, we reveal three novel aspects of how this is achieved. First, we show that the pool of nuclear, thus unphosphorylated YAP is approximately equally partitioned into daughter cells during cleavage divisions. This results in significant reduction of total YAP in each daughter cell compared to its mother. Second, we find that within a few hours of division total nuclear YAP levels adjust to newly acquired position/polarity states. Surprisingly, we find that this adjustment mainly manifests in the import of additional YAP into the nucleus in outer (polar) cells, rather than exporting nuclear YAP in inner (apolar) cells. Finally, we show that cell cycle-associated nuclear volume changes, result in prominent spikes of nuclear YAP concentration after divisions in all cells, irrespective of their position. These spikes dissipate as nuclei expand with cell cycle progression. As the combined volumes of two daughter nuclei eventually exceed that of the mother cell’s nucleus, there is a net nuclear volume increase through cleavage stages [24], [40], leading to a decrease in nuclear YAP concentration. Together these passive modes of regulation, involving dilution through cleavage divisions and changes in concentration due to nuclear volume dynamics lead to the emergence of cells with low nuclear YAP concentration, with active Hippo signaling likely only serving to prevent the entry of additional YAP into the nucleus. Interestingly, these dynamics result in cells reaching the low YAP state in two waves: either at the 16- or the 32-cell stages once the nuclear YAP concentration spikes “die down”. These YAP dynamics might explain the timing of SOX2 expression, which we previously observed to initiate also in two waves during the second half of the 16- and 32- cell stages [28]. The extent to which Hippo activity could remove YAP from the nucleus in the absence of these passive mechanisms remains to be investigated.

Previous work showed that while contractility differences among blastomeres did not directly correlate with YAP localization [16], [17], treatment with blebbistatin resulted in decreased nuclear YAP localization in polar cells [16]. Here we tested whether this observation was due to cell shape deformations, which can arise from contractility differences among cells or from blastocoel cavity formation. Surprisingly, we found that YAP was not responsive to mechanical perturbations that lead to cell shape deformations. This raises the possibility that, while not a factor mediating differential YAP localization among polar and apolar cells, contractility could be required in polar cells for nuclear YAP localization. It will be interesting to test whether contractility is required for Hippo signaling inactivation, and if so, identify the molecular mechanisms that are dependent on it.

## Acknowledgements

We thank Stas Shvartsman (Princeton University) for helpful conversations and scientific guidance. We thank Abhishek Biswas (Princeton University) for computational support, as well as Gary Laevsky and Sha Wang (Confocal Imaging Facility, Princeton University) for confocal microscopy support. Thank you to Bin Gu (Michigan State University) for sharing the YAP-miRFP670 transgenic mouse line and to Bradley Joyce (Princeton University) for embryo microinjection. This work utilized computational resources at Princeton University (Research Computing at Princeton University) and the Flatiron Institute (Scientific Computing Core). The Flatiron Institute is a division of the Simons Foundation.

## Contributions

Conceptualization: E.P., M.C., M.A. Methodology: M.C., M.A. Experimental data collection: M.C. Data processing and tracking: M.C. Computational modeling and analysis: M.A. Writing - original draft: E.P., M.C., M.A. Writing - review & editing: M.C., M.A., E.P. Supervision: E.P. Project administration: E.P. Funding acquisition: E.P.

---

**Supplemental Figure S1:**
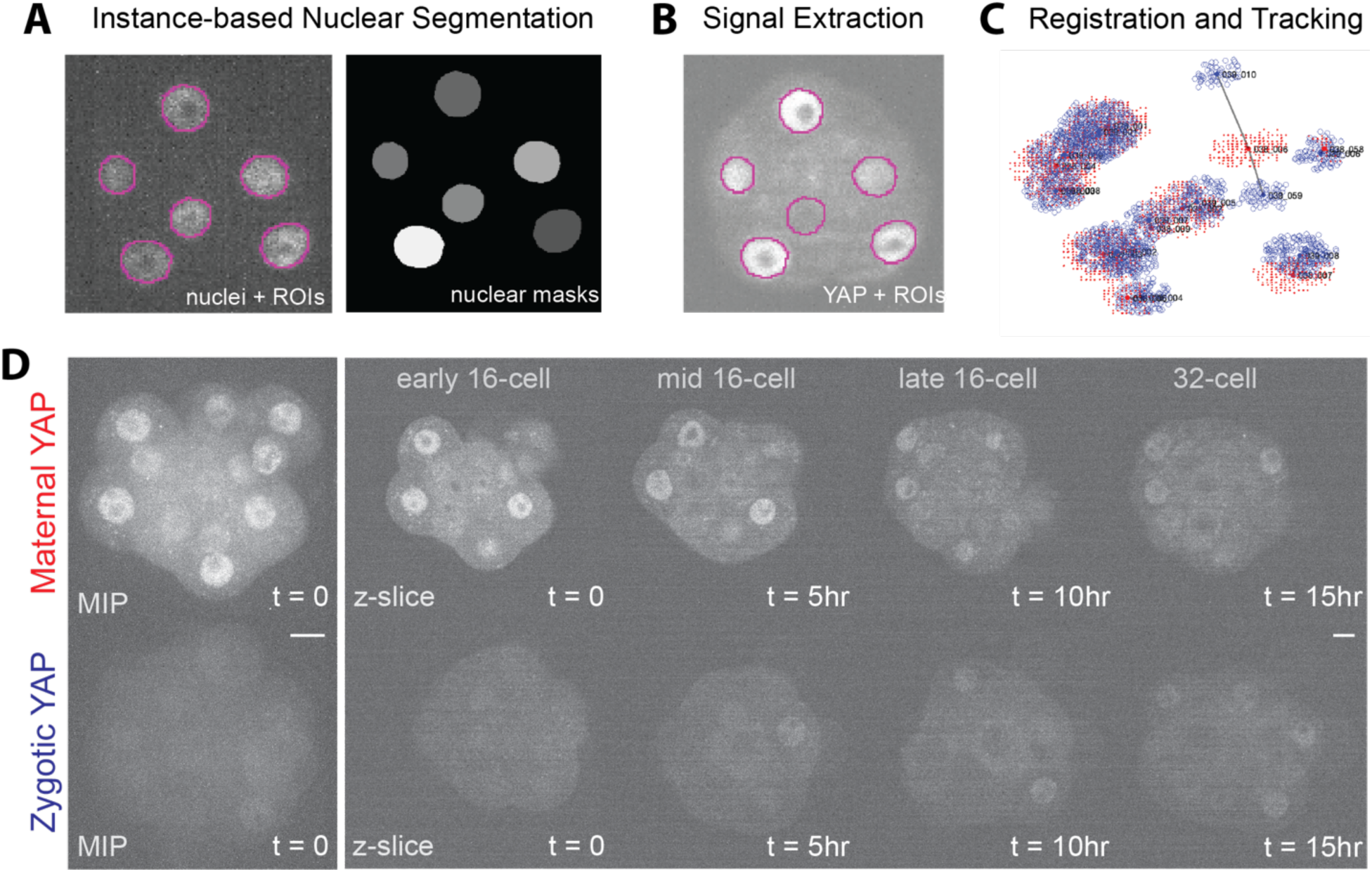
Image analysis method and maternal vs. zygotic YAP. A. Images illustrating the 3D-Stardist Instance-based nuclear segmentation method used to segment the nuclear channel. Regions of interest (ROI’s) are automatically generated from raw nuclear images and hand corrected before generating nuclear masks to be used for downstream registration and tracking. B. Nuclear YAP signal is extracted by overlaying nuclear ROI’s onto the raw YAP- miRFP670 images. C. Nuclear masks are translated into point clouds and first registered between timepoints before being tracked through divisions using a semi-automated tracking algorithm. Red point clouds represent time t, blue point clouds represent time t+1. Gray lines between point clouds illustrate automatic division detection. D. Spinning disk confocal time lapse images comparing maternal vs zygotic YAP- miRFP670. The first image shown at t=0 is a maximum intensity projection (MIP); all other images are of a single z-slice. All embryos were imaged in the same imaging session. All images are thresholded using the same brightness contrast settings. Scale bar 10um.

**Supplemental Figure S2:**
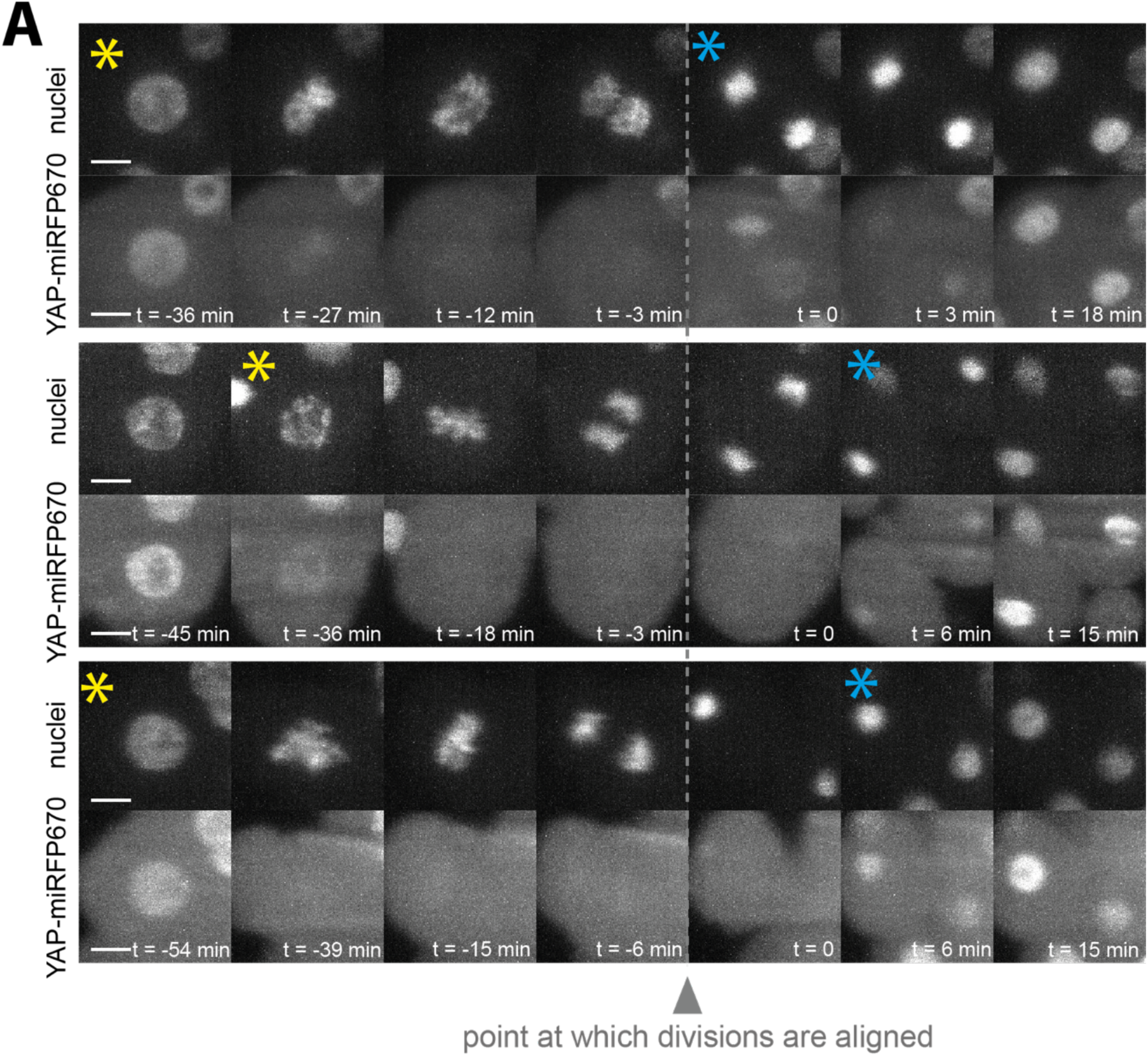
Detailed dynamics of nuclear YAP during mitosis. A. Image sequences of three different nuclei undergoing mitosis, demonstrating YAP-miRFP670 dynamics in relation to the time between nuclear envelope breakdown and reformation. Images were acquired every 3 minutes in this dataset. Dashed vertical line and arrowhead represent the time at which divisions are aligned for all analyses. Yellow asterisk: the last time point when the nuclear envelope is present. Blue asterisk: the first time point when the nucleus is reformed after mitosis. Scale bar 10um.

**Supplemental Figure S3:**
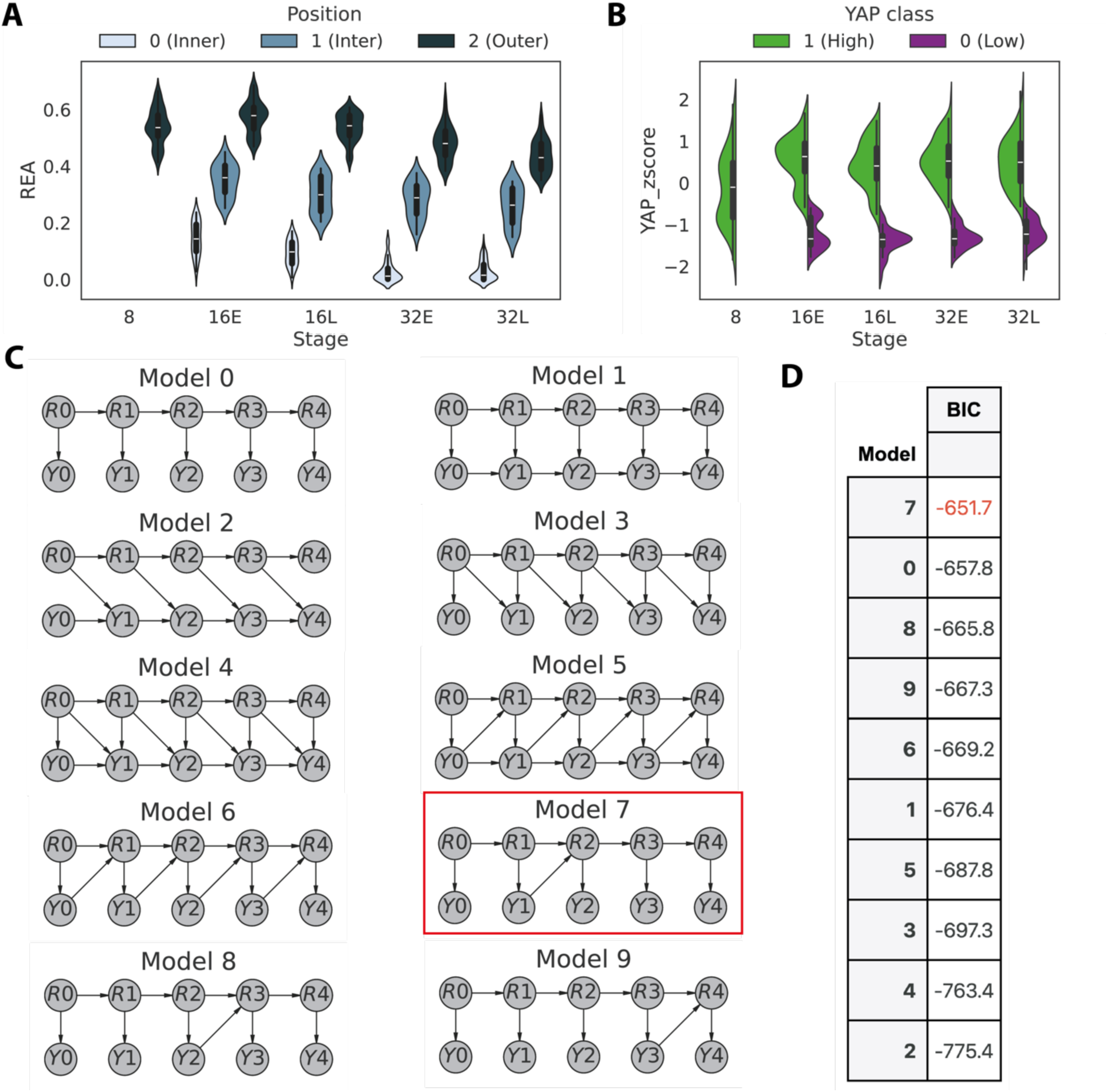
Bayesian model selection. A. Distribution of REA in inner, inter, and outer subpopulations at the 8-cell, early 16- cell, late 16-cell, early 32-cell, and late 32-cell stages. B. Distribution of nuclear YAP z-score in high- and low-YAP subpopulations at the 8- cell, early 16-cell, late 16-cell, early 32-cell, and late 32-cell stages. C. Ten model architectures tested for modeling REA-YAP dynamics. Model 7 was selected based on its Bayesian Information Criterion (BIC) score. D. Bayesian Information Criterion (BIC) for all models tested in panel (C), sorted from the highest (best) to the lowest (worst) BIC.

**Supplemental Figure S4:**
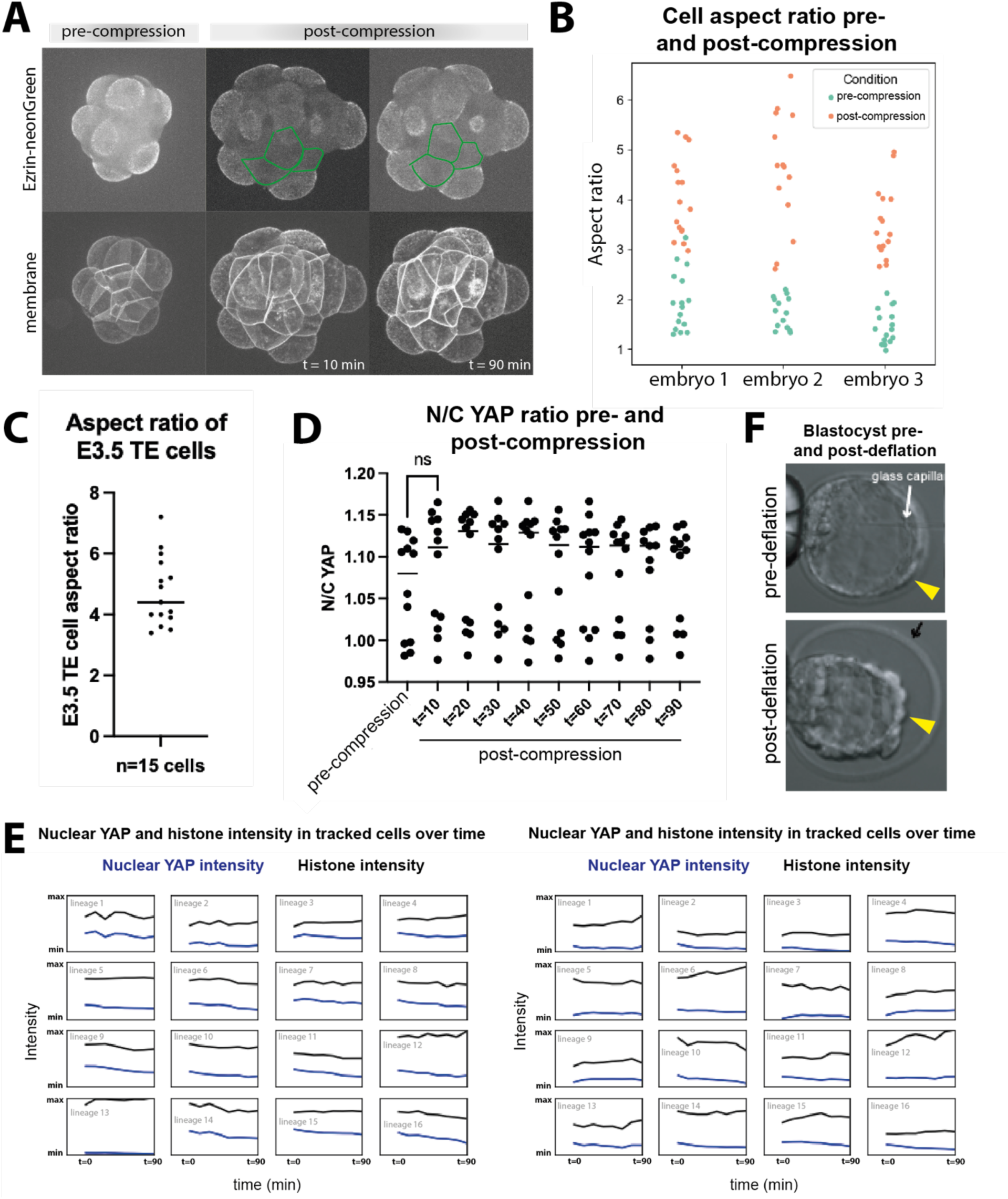
Mechanical perturbations of embryos. A. Confocal microscopy images of a 16-cell stage embryo expressing a membraneTomato and Ezrin-neonGreen pre- and post-compression. Green cell outlines highlight exposed cell surfaces that did not polarize during 90 minutes of compression. For all embryos imaged, compression either did not lead to cell rearrangements that significantly changed the exposed surface area of inner cells, or else exposed apolar cells at the onset of compression did not polarize during the imaging interval. N=5 embryos. B. A quantification of cell aspect ratio pre- and post-compression in three of the compressed embryos in Fig. 5C, demonstrating that cell aspect ratio increases as a result of compression. N=3 embryos. C. Aspect ratio of E3.5 TE cells as measured from n=15 cells sampled from 10 blastocysts, confirming that the aspect ratio of compressed cells in Fig. 5B is comparable to that of TE cells in blastocysts. D. In order to dissociate the effects of maternal YAP degradation from changes in nuclear YAP localization as a result of mechanical perturbation, nuclear:cytoplasmic (N/C) YAP ratios were measured for the compressed embryos in Fig. 3B and C. N/C YAP intensity was not statistically different between pre- and post-compression embryos, nor between post-compression values over the 90 minute time course. Shown is a quantification of N/C YAP pre- and post-compression for a single representative embryo. Student’s t-test results in no statistical difference between pre- and post-compression timepoints. E. Nuclei of compressed embryos analyzed in Fig. 5C were tracked through time. Nuclear YAP and histone intensities were extracted at each timepoint. Trends in nuclear YAP localization in single cells over time mirror trends in histone intensity, indicating that nuclear YAP levels do not increase in individual cell lineages over the course of compression (90min). Blue traces reflect nuclear YAP intensity; black traces represent histone intensity. The y-axis is scaled to the minimum and maximum of raw nuclear YAP and histone values within each embryo. Displayed are 16 tracked cell lineages from two sample embryos, representative of the behavior observed in N=9 compressed embryos. F. Example brightfield (BF) images of the method of blastocyst deflation. The top image shows a pre-deflation blastocyst being deflated with a fine glass capillary. The bottom image displays a post-deflation blastocyst, with a depressurized blastocoel cavity and rounded TE morphology. Yellow arrowheads point out examples of TE cells with a high aspect ratio (stretched) in the pre-deflation blastocyst and with low aspect ratio (rounded morphology) after blastocoel deflation.

## Materials and methods

### Transgenic mouse lines

Transgenic mouse lines for *Yap-miRFP670* (Gu 2020), *H2B- miRFP720* (Nunley 2024), and *mTmG* (Muzumdar 2007) mouse lines bred on a CD-1 (ICR) (Charles River) background were utilized in this study. All animal work was carried out at Princeton University, in accordance with the National Research Council’s *Guide for the Care and Use of Laboratory Animals*. Animal maintenance and husbandry were performed as per the guidelines outlined in the Laboratory Animal Welfare Act. All procedures were approved by Princeton University’s Institutional Animal Use and Care Committee (IACUC protocol #2133). Mice were cared for in a 14 hour daily light cycle facility maintained at 21°C and 48% average ambient humidity. See figure illustrations for genetic crosses used to derive embryos for each individual experimental purpose.

### Reagents for microinjection

*In vitro* transcription of mRNA performed as described previously (Gu 2018). Briefly, NotI (New England Biolabs, R3189L) digestion was used to linearize pCS2-H2B-miRFP720 plasmid (cloned into pCS2 vector using miRFP720 sequence obtained from Addgene plasmid #136560, Avdeeva et. al 2025) or pCS2-Cas9 plasmid (Addgene 122948) and an mMessage mMachine SP6 intro transcription kit (Thermo Fisher Scientific, AM1340) was used to synthesize the mRNA. mRNA was then purified using an RNeasy Cleanup kit (Qiagen, 74104) and eluted into RNAse-free water for aliquoting and storage at −80°C.

### Embryo isolation and microinjection

Embryos were isolated from natural mated females (6-15 week old) at either E1.5 (2-cell) or E2.5 (8-cell). Oviducts were flushed with M2 media (Zenith Biotech, M2116) and embryos were washed through microdrops of M2 under LifeGlobal paraffin oil (LGPO) (CopperSurgical, LGPO-500) before being transferred to precalibrated microdrops of KSOM EmbryoMax (Sigma, MR-101-D) under LGPO for culture in an incubator at 37°C, 5% CO_2_, 5% O_2_, 95% relative humidity. *H2B- miRFP720 mRNA* injection mixes were prepared in a nuclease-free injection buffer (10 mM Tris-HCl, pH 7.4 and 0.25 mM EDTA). E1.5 embryos were microinjected into both cells at the 2-cell stage with 75 ng/uL *H2B-miRFP720* mRNA, then cultured in KSOM under LGPO until experimental and/or imaging setup at E2.5.

### Light sheet microscopy

An inverted lightsheet microscope (InVi from Luxendo/Bruker) was used to acquire time lapse images. The microscope was outfitted with an incubation chamber maintained at a constant temperature of 37°C, 5% CO_2_, 5% O_2_, and 95% relative humidity throughout the duration of the imaging. A blunt tipped capillary was used to form ∼100um deep wells in the bottom of a v-shaped Fluorinated Ethylene Propylene (FEP) foil chamber (model # 80-0031-02-00). For each experiment, between eight to eighteen embryos are loaded into individual wells within the chamber. Z-stacks of each embryo are acquired sequentially in up to three consecutive channels every 15 minutes for a duration of approximately 30 hours. Images were acquired with 2.0um z-axis resolution and 0.208um x- and y-axis resolution. Fluorescent signal was captured using the following laser wavelengths and filters: mTmG 561nm excitation wavelength/577-612 BP emission filter; YAP-miRFP670 641nm excitation wavelength/659-690 BP emission filter; H2B- miRFP720 690nm excitation laser/700LP emission filter. All channels were acquired using the following Luxendo settings: 3.0 um beam expander, 10% beam intensity, and imaged on area mode. mTmG and H2B-miRFP720 channels were acquired at 150ms exposure; YAP-miRFP670 was acquired at 300ms exposure.

### Confocal microscopy

A spinning disk confocal microscope (W1, Nikon) equipped with an EMCCD camera (Hamamatsu) was used to acquire confocal time lapse and single timepoint images. For live image acquisition, embryos were placed into microdrops of KSOM EmbryoMax under LGPO on a MatTek dish (MatTek Corporation, P35G-1.5-20- C), and images were acquired using a 20X air objective with 2.8X magnification. For single timepoint images of fixed embryos, fixed embryos were imaged in PBS between two No. 1.5 glass coverslips using a 40X silicon oil objective at 1X magnification.

### Embryo compression

Embryos homozygous for Yap-miRFP670 and heterozygous for mTmG were isolated at E1.5 (2-cell) and injected with 50 ng/uL *H2B-RFP* mRNA into both blastomeres at the 2-cell stage. Embryos were cultured until E3.0 (16-cell), at which point they were transferred to individual microdrops in KSOM EmbryoMax under LGPO oil in a MatTek dish. Initial z-stacks of E3.0 embryos were acquired on a spinning disk confocal microscope (W1, Nikon). Embryos were then transferred to a new MatTek dish and compressed to a height of 30um between the glass bottom of the MatTek dish and a round glass coverslip, using 30um thick double sided tape as pillars (Nitto, PET-based double sided tape, No. 5603). The compressed embryos and coverslip were covered with LGPO to prevent evaporation during imaging. Embryo identity was maintained pre- and post-compression. Following compression, z-stacks were acquired of each embryo at 10 minute intervals for a duration of 90 minutes, using the same imaging settings as used to acquire pre-compression images.

### Blastocyst deflation

Embryos homozygous for Yap-miRFP670 and heterozygous for Cdx2-eGFP were isolated at E3.5 (blastocyst) and transferred to individual microdrops in KSOM EmbryoMax under LGPO oil on a MatTek dish. Initial z-stacks of E3.5 blastocysts were acquired on a spinning disk confocal microscope (W1, Nikon). Blastocysts were then transferred to a Leica DMi8 inverted microscope equipped with micromanipulators and deflated using a fine glass capillary. Cell-cell junctions in the mural TE were targeted for capillary insertion in order to minimize cell death. After deflation, embryos were transferred back to the MatTek dish for confocal imaging, and z-stacks of post-deflation embryos were acquired using the same imaging settings as for pre-deflation images. Embryo identity was maintained before and after deflation. Any TE cells that appeared lyzed following deflation (as judged by intense nuclear signal and compact morphology) were excluded from post-compression analysis.

### Image analysis of confocal microscopy data

Compressed embryo confocal microscopy data was analyzed by hand-segmenting nuclei in the nuclear channel (H2B- RFP670) to generate 3D nuclear ROI’s for all timepoints, encompassing initial pre-compression images through 90 minutes post-compression. Nuclear ROI’s were then tracked using the method described in Nunley et al. 2024, and transcription factor intensity was extracted from within the nuclear ROI’s for each point in time. Nuclear to cytoplasmic measurements were obtained by hand segmenting the central plane of the nucleus (excluding the nucleolus) and a corresponding section of the cytoplasm within the same plane for each cell. Transcription factor intensity was extracted from each ROI in Fiji, and N/C ratios were calculated by dividing nuclear signal by cytoplasmic signal for each cell over time.

Deflated blastocyst confocal microscopy data was analyzed by segmenting TE nuclei (using endogenously tagged CDX2-eGFP as a marker) employing the 3D-Stardist algorithm described in Nunley et al. 2024. The resulting TE nuclear ROI’s were checked and manually corrected in AnnotatorJ. YAP-miRFP670 intensity was extracted from the TE nuclei ROI’s for embryos pre- and post-compression.

### Analysis of light sheet data

Preprocessing and intensity extraction was applied to light sheet imaging data in accordance to [24], [28]. For every trajectory, the extracted signal was smoothed over a window of 2.5h duration within every cell cycle.

### Time warping

We normalize time for the length of the cleavage cycles at the level of individual branches. More precisely, the time within every cycle was first minmax normalized between 0 and 1 and rounded to the nearest multiple of 0.05 forming 20 warped timepoints per cycle. When more than one timepoint mapped to the same warped time, the average value was used. Warped times were then shifted according to the corresponding cleavage cycle, with [0,1], [1,2] and [2,3] corresponding to the 8, 16 and 32 cell stages respectively.

### Classification details

To classify branches by their YAP or REA status in Fig. 2 and 4, we normalized for the length of the cleavage cycles by dividing the cell cycle for every branch into “early” and “late” half stages of equal duration. More precisely, we used warped time to define early and late cycle stages by applying {x} < 0.5 and {x} >= 0.5 cutoffs on the warped times. We did not apply this cutoff to the 8-cell stage. As a result, 5 stages were defined: 8-cell, early/late 16-cell and early/late 32-cell stages. Before classification, first 2 hours post division were excluded from every branch, to normalize for small perturbations introduced by cell divisions.

### REA classification

REA measurements were obtained using methods from Porespy Python package. To classify cells as inner, intermediate and outer, REA was first averaged over every coarse-grained stage. We then applied k-means clustering on the resulting distribution for every stage using k=3, annotating the bottom (lowest mean) cluster as inner (0), intermediate cluster as intermediate (1) and top cluster as outer (2) cells (Fig. S3A).

### YAP classification

Classification of cells by their YAP status has been applied analogously to [28]. Briefly, in every embryo, YAP nuclear concentration was first z-scored at every timepoint, to normalize for the downward trends in its average nuclear concentration, and then averaged over every coarse-grained stage. Distributions of these averages were analyzed separately for every stage (Fig. S3B). At stages 16E and 16L, to separate the modes into low (0) and high (1), we extracted a kernel density estimate for each distribution using distplot method from the seaborn Python package, with default parameters, and identified its local minimum. For stages 32E and 32L, with no obvious bimodal structure present, we applied Gaussian mixture clustering with 3 clusters. Top two clusters, corresponding to intermediate and high YAP cells, were annotated as 1 while the bottom cluster was annotated as 0. Thus, individual thresholds were applied to each half stage; θ_1 = −0.59 for 16E, θ_2 = −0.82 for 16L, θ_3 = −0.61 for 32E, and θ_4 = −0.58 for 32L.

### Bayesian modeling

For dynamic Bayesian network (DBN) analysis (Fig. 4), we apply the approach that we established in [28]. Briefly, a non-homogeneous DBN was fit to (REA, YAP) pairwise dataset for every candidate structure (Fig. S3C) using methods from pgmpy Python package. Default parameters were used. Structure scores (BIC) were obtained using structure_score method with default parameters and compared. The winning structure shown in Fig. 4A was trained on the data with default parameters, i.e. using maximum likelihood approach. Fig. 4B, C, and D demonstrate the conditional probability distributions of the trained model, i.e., distributions of the form p(v|P(v)) for a vertex v and its parents, P(v). In Fig. 4C, the Y_1 node was marginalized out from p(R_2|R_1, Y_1).

### Membrane segmentation and analysis

To segment cells using the membrane channel of our imaging, we applied a pre-trained Cellpose model to the cropped images (www.cellpose.org) [27]. The model was applied using Python via command line with parameters “--pretrained_model cyto2 --do_3D”. A cutoff of 3000 pixels was then applied to every segment. The segmented nuclear images were then mapped to the segmented membrane images to find corresponding cell for every nucleus. More precisely, at every timepoint, every nucleus after segmentation and correction was mapped to the cell segment with the highest intersection. For robustness purposes, if several nuclei were mapped to the same cell, only the nucleus with the highest intersection over union was mapped.

### Image processing for figures and videos

Maximum intensity projections (MIPs) or single plane z-slices presented in all figures were processed in ImageJ by cropping the field of view to encompass the embryo and adjusting brightness contrast settings to obtain optimal visual contrast. Scale bars and timestamps were added in ImageJ. All analysis was carried out on raw data: images and videos were solely adjusted for visualization purposes.

